# An in-silico study of the mutation-associated effects on the spike protein of SARS-CoV-2, Omicron variant

**DOI:** 10.1101/2022.02.21.481269

**Authors:** Tushar Ahmed Shishir, Taslimun Jannat, Iftekhar Bin Naser

## Abstract

The emergence of Omicron (B.1.1.529), a new Variant of Concern in the COVID-19 outbreak, while accompanied by the ongoing Delta variant infection, has once again fueled fears of a new infection wave and global health concern. In the Omicron variant, the receptor-binding domain (RBD) of its spike glycoprotein is heavily mutated, a feature critical for the transmission rate of the virus by interacting with hACE2. In this study, we used a combination of conventional and advanced neural network-based in silico approaches to predict how these mutations would affect the spike protein. The results demonstrated a decrease in the electrostatic potentials of residues corresponding to receptor recognition sites, an increase in the alkalinity of the protein, a change in hydrophobicity, variations in functional residues, and an increase in the percentage of alpha-helix structure. Our next step was to predict the structural changes of the spike protein using the AI-based tool Alphafold2 and dock it with hACE2. The results revealed that the RBD of the Omicron variant had a higher affinity than the reference. Moreover, all-atom molecular dynamics simulations concluded that the RBD of the Omicron variant exhibits a more dispersed interaction network since mutations resulted in an increased number of hydrophobic interactions and hydrogen bonds with hACE2 compared to the reference RBD. In summary, our current study highlighted the potential structural basis for the enhanced transmissibility and pathogenicity of the Omicron variant, although further research is needed to investigate its epidemiological and biological implications.

## Introduction

COVID-19, caused by the SARS-CoV-2 virus, rapidly spread throughout the world and was declared a global pandemic as a public health emergency of international concern, which continues to have serious negative effects on health and the economy worldwide (Ozili and Arun, 2020; Wu et al., 2020). In order to halt the spread of SARS-CoV-2 viruses, several vaccines have been developed and implemented, but some of these efforts have been hindered by mutations leading to new virus variants (Mathieu et al., 2021; Tregoning et al., 2021; Wang et al., 2021). A recently emerged and rapidly spreading variant of SARS-CoV-2, named Omicron, has triggered worldwide concern, and the World Health Organization declared it a ‘variant of concern’ on November 26, 2021 (Dyer, 2021; WHO, 2021). A SARS-CoV-2 virus variant carrying mutations in its spike protein receptor-binding domain and increasing binding affinity in the RBD-hACE2 complex while speeding up the spread into human populations is known as a VOC (Variant of Concern) (Choi and Smith, 2021). A total of 30 mutations have been identified on the spike glycoprotein of the Omicron variant, of which 15 are located on its receptor-binding domain, and some of these mutations are also present in other VOCs, which may facilitate disease transmission, the severity of the disease, immune escape, and escape from diagnosis (Callaway, 2021; Karim and Karim, 2021; WHO, 2021).

A SARS-coV-2 infection begins with the binding of the viral spike protein to the Angiotensin-converting enzyme 2 (ACE2), which is then followed by the proteolytic processing of the trimeric spike protein (Scialo et al., 2020; Li et al., 2021). Mutations in the viral spike protein, therefore, play a very important role in viral evolution since these mutations affect SARS-CoV-2’s infectious properties and transmissibility and have a detrimental effect on vaccines and therapies (Harvey et al., 2021; Prévost and Finzi, 2021). Our present study uses an in silico approach to comprehensively analyze the spike protein of the Omicron variant with respect to the reference variant to distinguish the differences from a physicochemical, structural and functional perspective. Aside from this, the structure of the spike protein for the variant of the Omicron that has a number of mutations is not yet available; therefore, we used Alphafold2, a cutting-edge technology leading to structural predictions followed by a protein-protein docking of the ACE2 receptor in order to investigate the effect of mutations on protein interactions. Finally, we performed atomistic molecular dynamics simulations of the RBD-ACE2 complex of reference and Omicron variants and observed that the RBD of Omicron exhibits even stronger binding to hACE2. Molecular docking and MD simulation indicate that the stronger binding interactions may make that variant more infectious and more transmissible, and the current prevalence of this variant supports this assumption.

## Methods

### Data retrieval and annotation

SARS-CoV-2 reference spike protein sequences were obtained from Uniport with accession number P0DTC2 and the first complete Omicron genome from GISAID with accession number EPI_ISL_6640916 (GISAID, 2020; Bateman et al., 2021). The Omicron genome sequence was then annotated using the Cov-Seq program, followed by translation using the EMBOSS transeq tool (Liu et al., 2020)(Rice et al., 2000). A pairwise alignment of spike protein sequences was later performed with Clustal Omega, followed by the analysis of mutations (Sievers and Higgins, 2018). Furthermore, the PDB structure of the RBD-ACE2 complex with accession 6M0J and the PDB structure of the whole spike protein with accession 7N1U were used as references (Burley et al., 2019).

### Physicochemical parameter analysis

The spike protein of the reference and Omicron variants were subjected to preliminary sequence analyses to distinguish their physicochemical differences. The amino acid composition, molecular weight, distribution of charged residues, hydropathicity, aliphatic index, instability index, and a few other parameters are calculated mainly using the online server EMBOSS Pepstats (Rice et al., 2000). Further verification of the results from Pepstats was conducted using ProtParam, Prosite and AA-prop (Artimo et al., 2012; Bonnal et al., 2012). In addition, another web server called VOLPES was used to compare and visualize residue-level physicochemical properties (Bartonek and Zagrovic, 2019).

### Structural properties analysis

JPred4 was used to predict the secondary structure of spike proteins based on the JNett algorithm, which is one of the most advanced and accurate methods (Drozdetskiy et al., 2015). NetSurfP-2.0 tool was used to evaluate further the prediction, which utilizes convolutional and long short-term memory neural networks to predict secondary structure, solvent accessibility, and residual disorder (Klausen et al., 2019). Furthermore, the flDPnn server was used to predict intrinsically disordered regions and their functions in conjunction with NetSurfP-2.0 (Hu et al., 2021). Following that, the PredyFlexy server was used to determine the flexibility of the structures, while consurf and predictprotein were used to predict conserved regions (De Brevern et al., 2012; Ashkenazy et al., 2016; Bernhofer et al., 2021). Finally, AlphaFold2 was used to predict the tertiary structure of the Omicron spike protein (Jumper et al., 2021). Following the prediction of 3D structure, we calculated the RMSD and TM-score between the predicted Omicron spike protein and our reference spike protein (7N1U) using the TM-align tool and calculated the overlap of common contacts using the CMView program (Zhang and Skolnick, 2005; Vehlow et al., 2011).

### Functional properties analysis

In order to determine the effect of mutations on protein stability, Dyanmut2 and DeepDDG were used, where Dyanmut2 used normal mode analysis and graph representations of protein structures, and DeepDDG used neural networks to predict the effect of mutations on protein stability (Cao et al., 2019; Rodrigues et al., 2021). In addition, several tools, including SNAP2, PROVEAN and SIFT, were used to assess the impact of mutations on function (Sim et al., 2012; Choi and Chan, 2015). As a final step, we predicted how mutations would affect the propensity of SARS-CoV-2 to cause disease using the VarSite webserver (Laskowski et al., 2020). Whenever more than one tool was employed to predict the effects, the common outcomes were taken into consideration.

### Molecular docking and protein-protein interaction analysis

First, we retrieved the RBD-ACE2 protein complex with accession number 6M0J and separated the chains. Then, the Pymol mutagenesis wizard was used to introduce the specific mutations at the appropriate residues in the receptor-binding domain. After preparing the protein, Cluspro and HDOCK were used to dock the reference and Omicron RBD to the ACE-2 receptor (6M0J, chain A) (Kozakov et al., 2017; Yan et al., 2020). We then used the PRODIGY webserver to calculate the binding affinity and the PIC server to investigate the interactions between RBD and hACE2 (Tina et al., 2007; Xue et al., 2016). Finally, the Pymol graphical software was utilized for figure generation (DeLano, 2020).

### Molecular dynamics simulation

All-atom MD simulation was carried out in GROMACS 2021.2 software package, and ACE2-RBD protein complexes of both reference and Omicron variants were prepared using the CHARMM36 force fields (Van Der Spoel et al., 2005; Yu and Klauda, 2020). Each protein complex was then solvated with the TIP3P water model by adding 0.15 mM sodium chloride within a dodecahedron box. The distance between the protein complex and the corner of the box was set to 1.2 nm. The system energy was minimized with the Steepest Descent algorithm in 50,000 steps, followed by the system was equilibration in two phases. Firstly, 10ps NVT (constant number of particles, volume, and temperature) simulation was performed to equilibrate the temperature at 310.15 K guided by V-rescale temperature coupling algorithm, followed by 100ps NPT (constant number of particles, pressure, and temperature) simulation to equilibrate the system at 1 atm pressure and 310.15 K by using Parrinello-Rahman barostat algorithm (Martoňák et al., 2003). Finally, the MD simulations of both reference and Omicron variant ACE2-RBD systems were run for 200ns with a time step of 2.0 fs under NPT ensemble using GROMACS 2021.2 software and long-range electrostatic interactions were computed using Particle Mesh Ewald (PME) algorithm. The cutoff values of the electrostatic and Van der Waals interactions were set to 12 Å while the linear constraint LINCS algorithm was used to constrain all covalent bond lengths, including hydrogens.

MD trajectories were analyzed using GROMACS’ integrated tools for computing root-mean-square deviation (RMSD) and difference root-mean-square fluctuation (RMSF). Lastly, we investigated hydrogen bond interactions and their relative frequencies with the VMD package, setting the hydrogen bond distance and angle to 3.0 Å and 20°, respectively, and calculated the binding energy using the Prime 3.0 MM-GBSA module (Humphrey et al., 1996).

## Results

SARS-CoV-2 Omicron variant has been designated as the variant of concern due to its rapid emergence worldwide, which includes 30 mutations in the Spike protein, and nearly half of them are in the receptor-binding domain (Fig 1). Due to sequence loss, the Omicron variant has 1270 amino acids instead of the reference spike’s 1273 amino acids. It is evident from a primary analysis of the protein sequence that this variant has more Arginine, Histidine, Lysine and Glutamic acid than the reference, indicating that the spike protein is more charged (Supplementary file). Furthermore, these residues are exposed to a much greater extent and contribute to binding with receptors because their pKa’s are high enough with polar side chains, which can form hydrogen bonds. On the other hand, Isoleucine and Phenylalanine are also present in higher numbers within the protein’s core, making the spike protein more hydrophobic than the reference variant. These mutations will alter its physicochemical and structural properties, which will affect the transmission rate and pathogenicity within human populations by reducing antibody-mediated protection.

**Fig 1.**
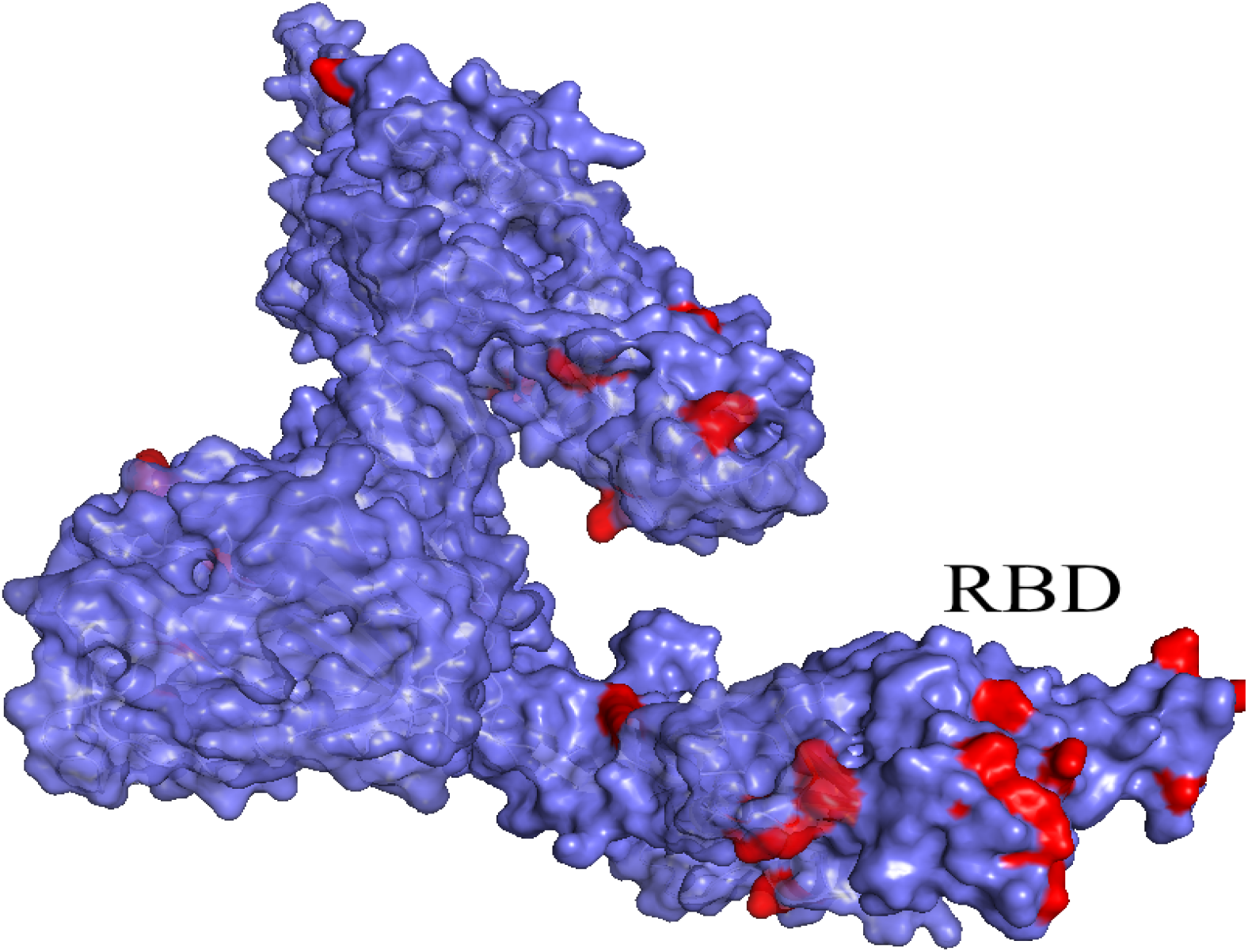
Three-dimensional structure of the Omicron variant spike protein monomer. Mutations are highlighted in red, and nearly half of them occur in the receptor-binding domain.

### 3.1 Effects on physicochemical properties

Despite having three fewer amino acids, the molecular weight of the Omicron variant (141328.11 Da) is higher than the reference variant of (141178.47 Da), and the mutations are biased toward the nonpolar amino acids (Fig 2A); therefore, the hydrophobicity of the spike protein of the Omicron variant increased (Table 1 and Fig 2B). While the number of hydrophobic residues increased, the GRAVY (Grand Average of Hydrophilicity) value indicates the protein has become slightly more hydrophilic intrinsically, which is indicative of the effects of mutations on the surface accessibility of the protein (Fig 2C) due to alteration of the secondary and tertiary structural properties. Furthermore, table 1 shows an increment of both acidic and basic residues in this variant; however, the increase in basic residues is higher (Fig 2D), resulting in a net charge of 8, which is likely to facilitate the interaction with hACE2. There was less electronegativity observed among the residues closest to the recognition of the receptor-binding domain of spike protein where the ACE-2 receptor binds (Fig 3C and 3D). In contrast, a high level of electronegativity was evident among the other residues of the domain and the complete protein of the Omicron variant (3A and 3B).

**Table 1.** Differences in the physicochemical properties of reference and Omicron variant spike proteins.

| Variant | Molecular weight | Polarity |  | GRAVY | Charged Residues |  | Net Charge | Isoelectric Point | Extinction coefficient (1mg/ml) | Instability index | Aliphatic index |
| --- | --- | --- | --- | --- | --- | --- | --- | --- | --- | --- | --- |
|  |  | Polar | Non-polar |  | Acidic | Basic |  |  |  |  |  |
| Reference | 141178.5 | 45.25% | 54.75% | -0.079 | 8.64% | 9.42% | 1.5 | 6.62 | 1.037 | 33.01 | 84.67 |
| Omicron | 141314.04 | 45.12% | 54.88% | -0.080 | 8.74% | 10.08% | 8.0 | 7.18 | 1.036 | 34.57 | 84.95 |

**Fig 2.**
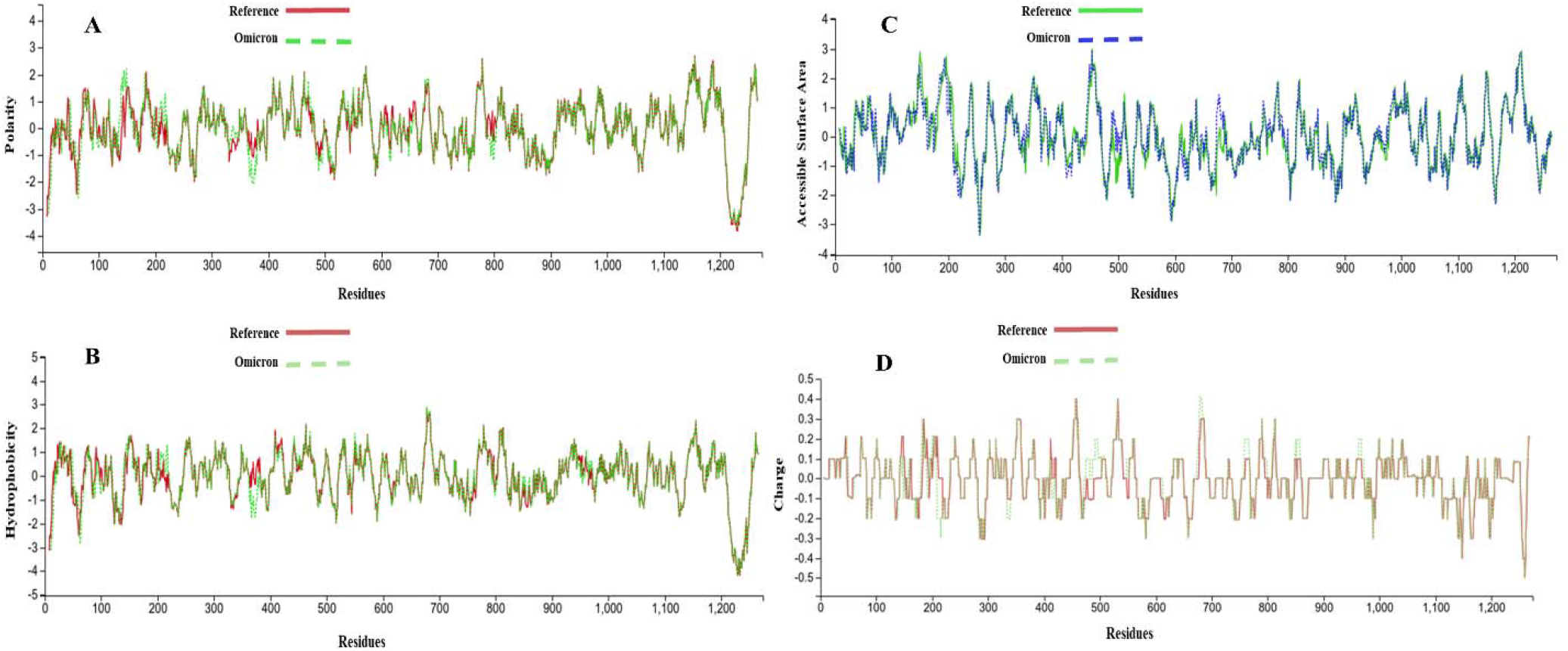
Physicochemical properties of reference variant and Omicron variant spike proteins at the residue level. In every parameter, the receptor binding domain region shows larger fluctuations. **A**. The difference in polarity, **B**. Differences in hydrophobicity, **C**. Variations in the accessible surface area, **D**. Variations in electrostatic potential

**Fig 3.**
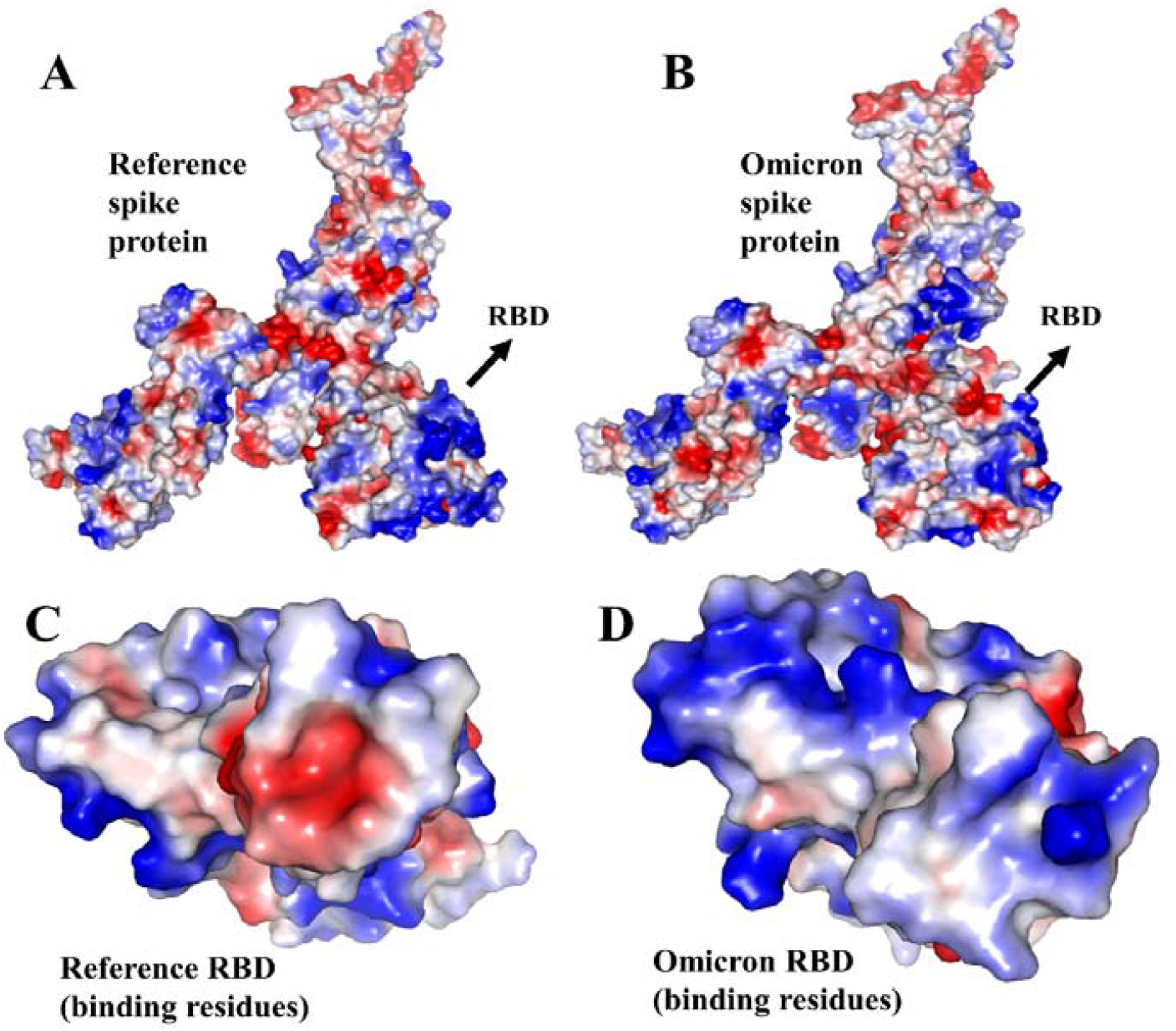
The electrostatic surface of the spike protein. The red and blue colors denote negative and positive potential respectively. Due to mutations at residues near the receptor recognition site, their electrostatic potential shifts from negative to positive, predicted to facilitate the binding with hACE2. **A**. Electrostatic potential map for the whole reference spike protein, **B**. Overall electrostatic potential map of the whole Omicron variant spike protein, C. High electronegativity of the recognition site at RBD of reference variant, **D**. Low electronegativity of the recognition site at RBD of Omicron variant.

Moreover, the Omicron variant’s isoelectric point (pI) is 7.18, meaning the protein is slightly alkaline, whereas the reference variant is acidic in nature with a pI value of 6.62. Then, another important physicochemical parameter is the extinction coefficient, which is the measure of how much light is absorbed by the polypeptides. The extinction coefficient of the Omicron variant was calculated to be 1.036, while 1.037 was that of the reference variant assuming all cysteine residues are reduced. Moreover, we found that both variants of the spike proteins were stable with scores of 33.01 and 34.57 respectively for reference and Omicron, with reference being more stable. Finally, both the reference and Omicron variants had higher aliphatic index values of 84.67 and 84.95, respectively, indicating that both variants are thermostable over a wide temperature range.

### 3.2 Effect on structural properties

Mutations in the spike protein were predicted to affect its structural properties. First of all, according to the secondary structural analysis, this variant has a higher fraction of alpha-helix (23.46%) than the reference (21.52%), while the beta-strand structure was decreased (Table 2). The T470-Q474 residues of the receptor-binding domain transitioning into the alpha-helix structure would increase the RBD’s stability, making the variant more transmissible and pathogenic since hACE2 interacts with the T470-F490 loop. Overall, ten residues of beta-strands and coils were predicted to be transformed into alpha-helix, but the opposite was not observed. There were, however, fourteen beta-strand residues predicted to be transformed into random coils, while seven random coil residues may be transformed into beta-strand residues (Supplementary file). Then, the mutations influenced the solvent accessibility of 154 residues and made the variant more hydrophilic because a higher number of residues were exposed. Among 154 residues, 61 were exposed from the buried or intermediate state, while 54 were buried (Supplementary file).

**Table 2.** Differences in structural features between reference and Omicron variant spike proteins.

| Variant | Secondary Structure |  |  | Solvent accessibility |  |  | Residual flexibility |  |  | Conserved Residue |  |
| --- | --- | --- | --- | --- | --- | --- | --- | --- | --- | --- | --- |
|  | Alpha Helix | Beta Strand | Coil | Exposed | Intermediate | Buried | Flexible | Intermediate | Rigid | Functional | Structural |
| Reference | 21.52% | 22.07% | 56.40% | 30.48% | 9.19% | 60.33% | 361 | 588 | 304 | 321 | 354 |
| Omicron | 23.46% | 20.55% | 55.98% | 31.42% | 8.66% | 59.92% | 353 | 586 | 311 | 309 | 394 |

Additionally, mutations in the spike protein changed the residual flexibility and increased the rigidity of the protein, which would affect its functionality. In the reference variant, 361 residues were predicted to be flexible, but the number decreased to 353 while rigid residues increased from 304 to 311 (Table 2). While some flexible residues gained an intermediate state without becoming rigid, very few rigid residues developed a flexible state directly, and all rigid transitions occurred among residues of the intermediate state (Supplementary file). In this variant, flexibility predictions showed that the transmembrane domain and heptapeptide repeats (1213-1237, 912–984 and 1163–1213 residues) of the S2 subunit are highly flexible, which could affect the viral cell fusion with host cells. Despite the mutations affecting the protein’s residual flexibility, we did not observe any significant changes in the residual disorder of the protein implying several algorithms. Finally, the Omicron variant of the spike protein has 703 conserved residues for structure and function, compared to 675 residues in the reference variant (Table 2 and supplementary file). It was found that the structural residues of this variant increased from 354 to 394, which would likely increase the stability of the protein. There was, however, a decrease from 321 to 309 functional residues, which may have an impact on the viral fatality.

In addition, both protein structure and conformation dynamics are associated with biological functions, so we analyzed the reference and the Omicron variant’s tertiary structure further to find the structural variations. We observed an RMSD of 0.20 and a TM-score of 0.99780 (Fig 3A) between the spike proteins, indicating a higher degree of structural similarity. In contrast, a contact map overlapping analysis yielded 90.5% common contacts, with the reference variant having 89 unique contacts among the residues, whereas the Omicron variant had 438 (Fig 4A). The contact map analysis indicates some differences among the functional residues of the protein despite the RMSD value and TM-score indicating no major structural changes caused by the mutations.

**Fig 4.**
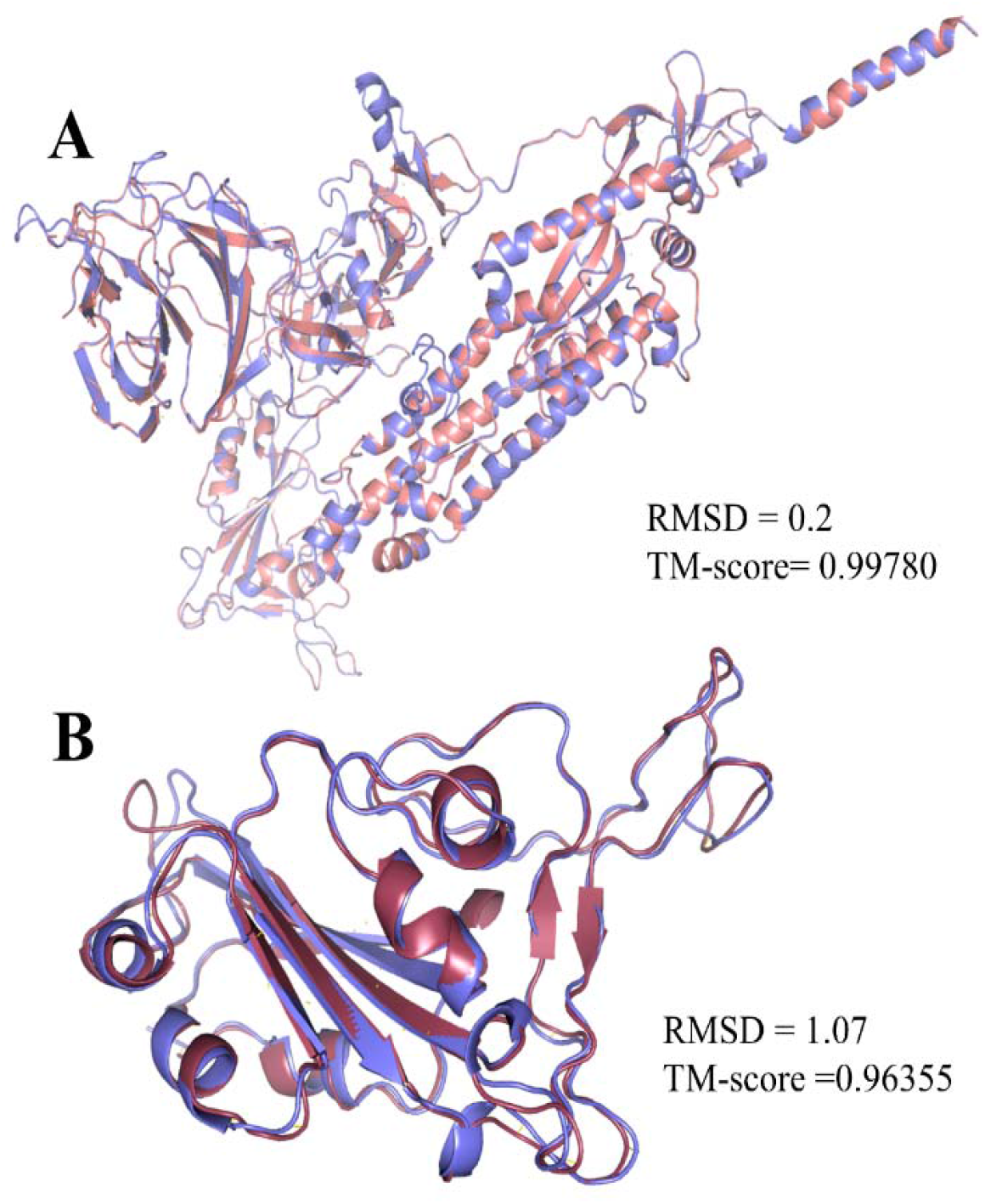
Comparison of the spike protein and the RBD of the reference and Omicron variants. In superimposing representative structures, no major changes are seen in the whole spike protein, but in the RBD, there is a difference in atomic coordinates, which would affect its interaction with receptor hACE2. **A**. Superimposed structure of the whole spike protein of the reference and the Omicron variant, **B**. Superimposed receptor-binding domain structure of the reference and the Omicron variant.

The superimposition of the receptor-binding domain structures yielded an RMSD value of 1.07, indicating higher structural differences in atomic coordinates than the reference variant; however, the TM-score of 0.96355 suggests no major structural differences occurred (Fig 3B). A total of 91.3% of residue contacts were common between the reference and Omicron variant, with 50 and 34 unique contacts respectively (Fig 4B), which indicates differences in the functional residues of the RBD that may affect viral transmissibility.

### 3.3 Effect on stability, functionality and disease propensity

Using a combination of deep learning neural network algorithms and structure-based predictions, the effect of the mutations on the spike protein stability was predicted. There is a decrease in structural stability for all amino acid changes, but the S142H, N764K and P681H are predicted to have a significant impact (Table 3). The functional effect analysis revealed that the E484A, Y505H, T547K, N764K, N856K, and N969K mutations impair the spike protein’s function, and the rest are neutral (Table 3). One of the five mutations, E484A, is located in the receptor-binding domain, so this mutation would likely influence viral transmission. The rest of the mutations in the RBD were predicted not to affect protein function but to reduce its structural stability, which could affect it either way.

**Table 3.** The effects of mutations on the stability, functionality, and disease-proneness of the Omicron variant.

| Reference AA | Position | Variant AA | Effect |  |  |
| --- | --- | --- | --- | --- | --- |
|  |  |  | Stability | Functionality | Disease propensity |
| A | 67 | V | Decrease Stability | Neutral | Less disease prone |
| T | 95 | I | Decrease Stability | Neutral | Less disease prone |
| G | 142 | D | Decrease Stability | Neutral | More disease prone |
| L | 212 | I | Decrease Stability | Neutral | Less disease prone |
| G | 339 | D | Decrease Stability | Neutral | More disease prone |
| S | 371 | L | Decrease Stability | Neutral | More disease prone |
| S | 373 | P | Decrease Stability | Neutral | More disease prone |
| S | 375 | F | Decrease Stability | Neutral | Less disease prone |
| K | 417 | N | Decrease Stability | Neutral | Less disease prone |
| G | 446 | S | Decrease Stability | Neutral | More disease prone |
| S | 477 | N | Decrease Stability | Neutral | Less disease prone |
| T | 478 | K | Decrease Stability | Neutral | More disease prone |
| E | 484 | A | Decrease Stability | Affect | Less disease prone |
| Q | 493 | R | Decrease Stability | Neutral | Less disease prone |
| G | 496 | S | Decrease Stability | Neutral | More disease prone |
| Q | 498 | R | Decrease Stability | Neutral | Less disease prone |
| N | 501 | Y | Decrease Stability | Neutral | More disease prone |
| Y | 505 | H | Decrease Stability | Affect | More disease prone |
| T | 547 | K | Decrease Stability | Affect | More disease prone |
| D | 614 | G | Decrease Stability | Neutral | Less disease prone |
| H | 655 | Y | Decrease Stability | Neutral | Less disease prone |
| N | 679 | K | Decrease Stability | Neutral | More disease prone |
| P | 681 | H | Decrease Stability | Neutral | Less disease prone |
| N | 764 | K | Decrease Stability | Affect | More disease prone |
| D | 796 | Y | Decrease Stability | Neutral | More disease prone |
| N | 856 | K | Decrease Stability | Affect | More disease prone |
| Q | 954 | H | Decrease Stability | Neutral | Less disease prone |
| N | 969 | K | Decrease Stability | Affect | More disease prone |
| L | 981 | F | Decrease Stability | Neutral | Less disease prone |

Additionally, sixteen mutations were predicted to increase disease propensity and thirteen to decrease it. Together with the other twelve mutations, it was predicted that the E484A mutation would decrease the probability of diseases induced by the protein, which was predicted to affect the protein function. On the other hand, the other four protein function impairing mutations are predicted to increase the likelihood of disease (Table 3).

### 3.4 Effect on binding interaction with hACE2

Infectivity, transmission, and pathogenesis are largely determined by the binding affinity of the virus towards the receptor, so mutations in the receptor-binding domain of the virus can greatly influence its activities. Since SARS-CoV-2 interacts with hACE2 through its C-terminal domain, mutations at key residues would affect the interaction with hACE2. In this part of the study, we found that the binding affinity of the Omicron variant differs from the reference variant. Our results from the HDOCK server showed that the docking score and binding affinity for the Omicron variant were -343.56 and -11.8 kcal/mol, compared to -310.19 and -11.5 kcal/mol for the reference variant (Table 4). Moreover, the ClusPro server provides us with the docking score and binding affinity for Omicron variants -703.8 and -14.7, respectively, compared to -639.3 and -11.9 for the reference variant (Table 4). Thus, it is evident that the Omicron variant exhibits a stronger binding affinity to the hACE2 than the reference variant, implying a potential for higher viral transmission.

**Table 4.** Differences in interfacial contacts between reference and Omicron variant with hACE2.

| Interfacial contacts |  | Reference variant | Omicron Variant |
| --- | --- | --- | --- |
| Binding affinity (kcal/mol) | 1. ClusPro | -11.9 | -14.7 |
|  | 2. HDock | -11.5 | -11.8 |
| Hydrophobic interaction |  | 4 | 6 |
| Main chain-main chain hydrogen bonds |  | 1 | 0 |
| Main chain-side chain hydrogen bonds |  | 5 | 14 |
| Side chain-side chain hydrogen bonds |  | 11 | 18 |
| Ionic interactions |  | 3 | 7 |
| Aromatic-Sulphur interactions |  | 1 | 0 |
| Pi-cation interactions |  | 0 | 3 |

There are significantly more interactions with hACE2 than in the reference (Fig 5), with 48 versus 25 interfacial contacts in the Omicron variant, most of which are hydrogen bonds (Table 4). Our study of protein-protein interaction in the reference variant found four hydrophobic interactions involving Y489 and F486 residues. These interactions were not present in the Omicron variant; however, six new hydrophobic bonds involving Y446, K452, I469, A481 and F487 were discovered. Pi-Cation interactions were observed at three new sites with Y446, Y486 and Y498 residues in the Omicron variant, but there were no Pi-Cation interactions in the reference variant, while the only aromatic bond of the reference was absent in Omicron. Additionally, the number of ionic interactions increased from three to seven when the RBDs of Omicron spikes bind with hACE2. However, the maximum discrepancy was observed regarding hydrogen bond formation, where the only hydrogen bond between the main chains of the reference RBD and hACE2 was absent from the Omicron variant. Meanwhile, the number of bonds formed between the main chain and side chain increased dramatically from 5 to 14, and the number of bonds formed between side chains rose from 11 to 18.

**Fig 5.**
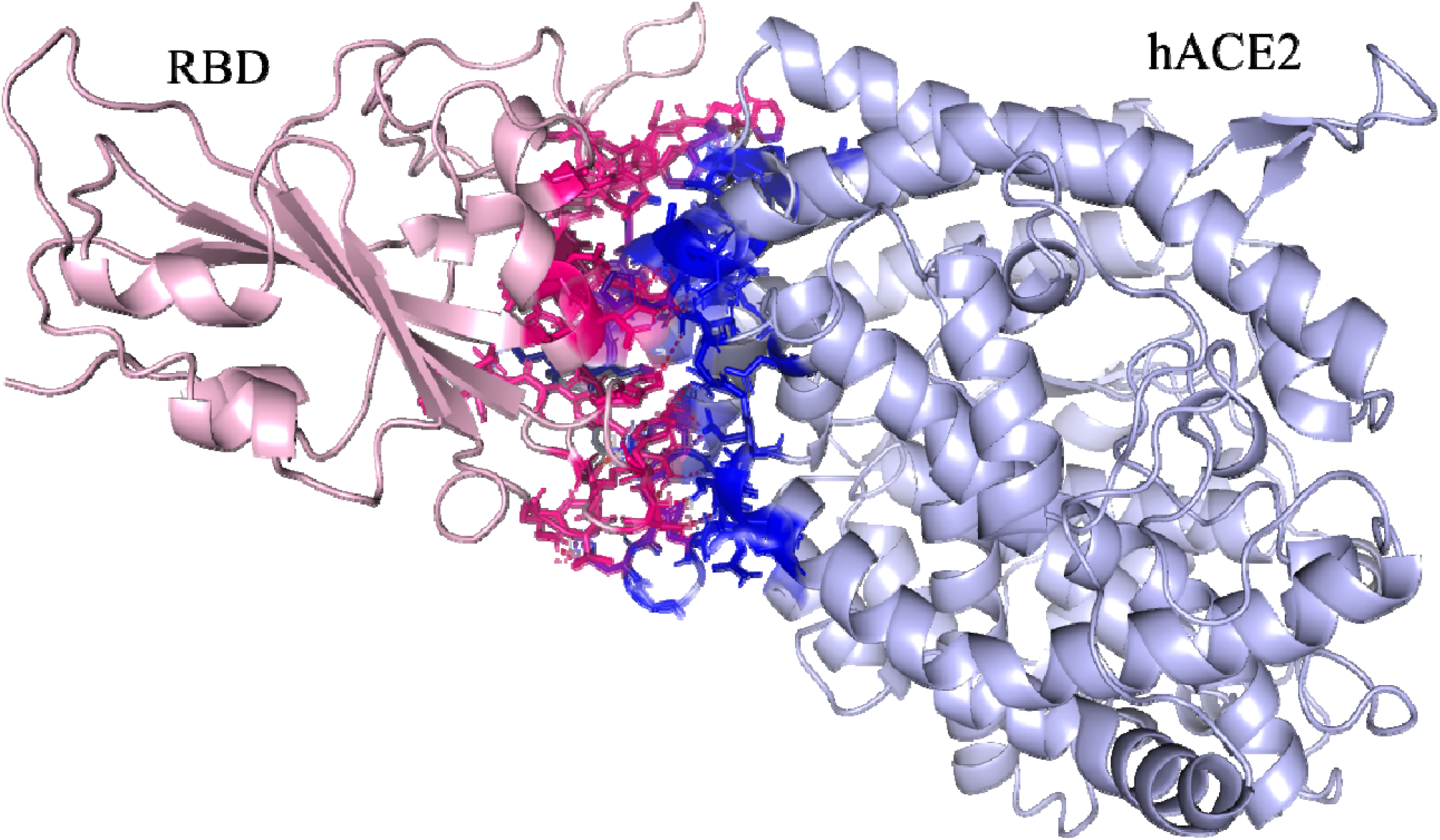
Interfacial contacts of the receptor-binding domain of the Omicron variant with hACE2.

Six of the fifteen mutations found in the RBD of the spike protein in the Omicron variant directly affected its interaction with hACE2. For instance, E484 was in ionic contact with K31 in th reference structure, but the Omicron variant (E484A) lost this ionic interaction; instead, A484 formed two hydrophobic interactions with L79 and M82 residues of hCAE2. The most drastic effect on the interactions took place due to the Q498R mutation, which is located near the hACE2 recognition loop structure of RBD. There was a hydrogen bond present between the Q498 of the reference RBD and the Q42 residue of hACE2; however, the Q489R mutation impaired that bond and led to the formation of several new hydrogen bonds among the residues of the main chains and side chains, as well as ion interactions with E208 and D216 residues of hACE2 (Supplementary file). In addition, the N501Y mutations, which were previously reported for the alpha variant, also increased the binding affinity for the Omicron variant because two hydrogen bonds and one Pi-Cation bond were formed instead of one hydrogen bond in the reference variant. In the supplementary file, we provide a full comparison of the protein-protein interactions of Omicron RBD and reference RBD with hACE2. To summarize, it is evident that mutations increased the binding affinity between the receptor-binding domain and hACE2, which is a key factor in the transmissibility and pathogenesis of SARS-CoV-2.

### 3.5 MD simulation result on RBD-ACE2 complex

Using molecular dynamics simulation, we observed new interactions formed with hACE2 while retaining many of the old ones found in the reference sequence. These interactions were attributed to additional residues at the interface (Q474, G476, N477, K478, A484, F486, and Y489). Despite this, we found several interaction disruptions; for instance, the salt bridge between E484 and K31 was lost, the hydrogen bond between K417 and D30 was damaged, and other interactions were disrupted at residues F456, A475, and G502. According to our analysis of 15 mutations that have been reported to occur in the RBD of the Omicron variant, 6 of these mutations were found to have additional effects on the binding of the Omicron RBD with hACE2.

In addition, on average, we found 6.89±1.28 hydrogen bonds between Omicron RBD and hACE2, as compared to 5.52±1.26 hydrogen bonds with the reference variant (Fig 6C). This indicates that the Omicron variant enhances the interaction between RBD and hACE2 compared to the reference variant. Interestingly, mutations of Q498R and N501Y were found to form new hydrogen bonds with occupancy higher than 15%. In contrast, mutations of K417N and Y505H led to the loss of hydrogen bonds, and some mutations did not affect hydrogen bonding, for example, the K493-E35 bond resulting from the Q493K mutation. After that, we calculated the binding energies of the hACE2 complex in comparison to either the RBD of Omicron and the reference, and it was found that the binding energy of hACE2 is lower when binding to Omicron RBD (−98.02 kcal/mol) as opposed to the reference (−83.7 kcal/mol).

**Fig 6.**
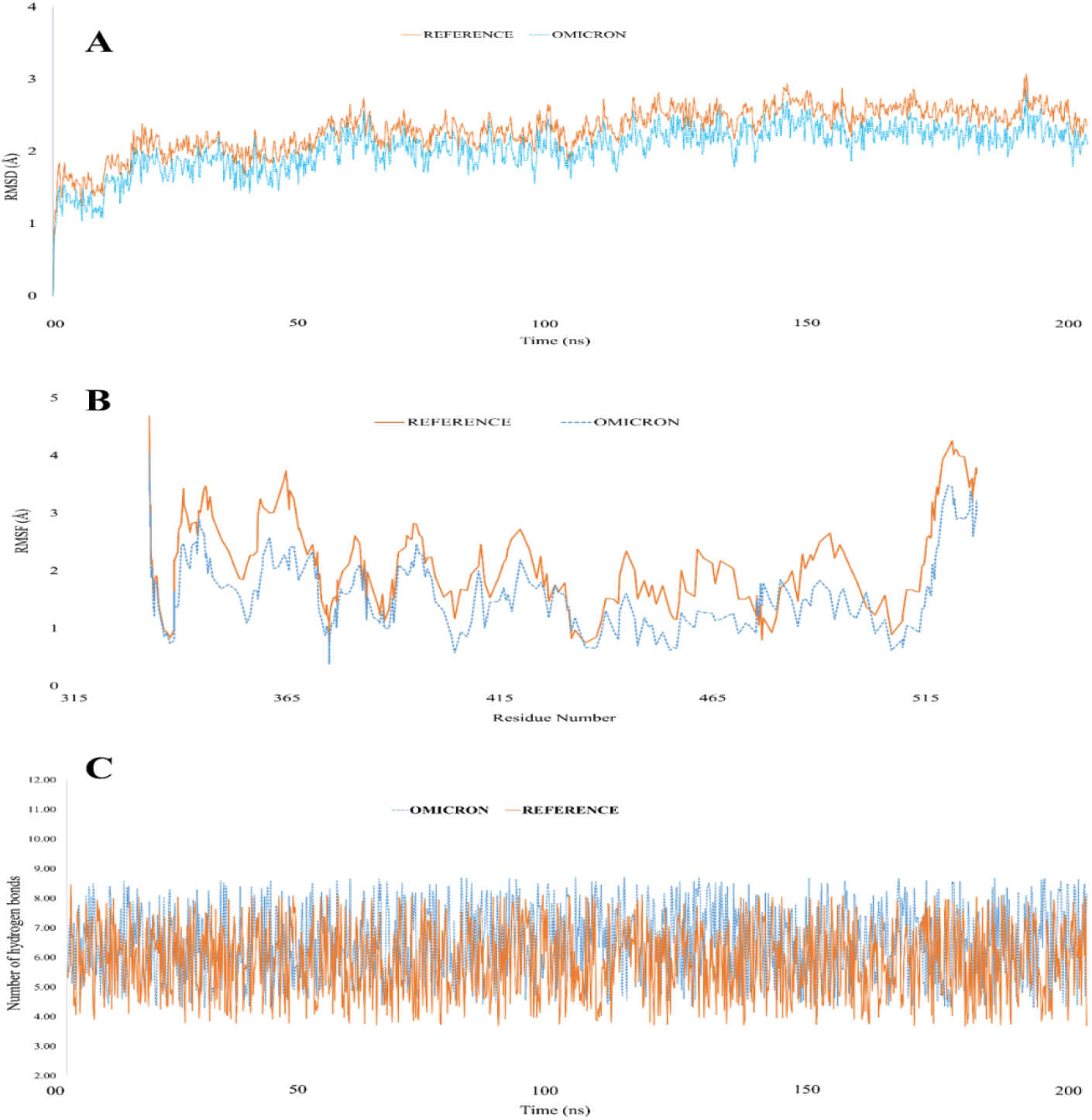
A representation of the stability and hydrogen bonds in the RBD-hACE2 complex structure in the reference and Omicron variants. **A**. Root mean square deviation (RMSD) of RBD-hACE2 complex. **B**. Root mean square fluctuations (RMSF) of the receptor binding residues in RBD-hACE2 complex. **C**. Number of hydrogen bonds of the RBD observed with hACE2 during simulations.

In addition, the root-mean-square-deviation (RMSD) for the reference ACE2-RBD complex was 2.61 Å (Fig 6A). In comparison, the value was 2.05 Å for the Omicron RBD-ACE2 complex; a finding that indicates the mutations did not lead to significantly reduced stability of RBD but instead increased the stability of the Omicron RBD-ACE2 complex over the reference variant. Additionally, we calculated the root-mean-square fluctuations (RMSF) from the trajectory data and found that the RBD of the Omicron variant is more rigid than the reference variant and that the RMSF values averaged to 1.7 Å and 2.2 Å, respectively (fig 6B). Interestingly, the mutated residues at the interface of the hACE2 showed reduced fluctuations compared to the reference variant at the interface. Thus, when taking into account the number of hydrogen bonds, binding energies, RMSD, RMSF, and RMSD, Omicron RBD-ACE2 appears to be more stable than the reference complex.

## Discussion

The World Health Organization deems Micron a variant of concern due to its rapid transmission rate and the unusually large number of amino acid mutations in its spike protein. In this study, we investigated how the mutations affect the spike protein from physicochemical, structural and functional standpoints and how they modulate its interactions with the host protein ACE2. To begin with, despite a decline in overall hydrophobicity resulting from changes in surface accessibility of several residues caused by the mutations in the Omicron variant of spike protein, the hydrophobic residues are increased in number, which are required for the stability of the protein and make the protein’s core (Fersht and Serrano, 1993; Matthews, 1993). In addition, changes in the hydrophobicity of amino acids may alter the structure of the epitope in the receptor-binding domain of the spike protein, which may affect the immune response to the virus (Tekewe et al., 2016). Moreover, we found that the protein was more alkaline than the reference and observed lower electronegativity in residues near the receptor recognition site.

In addition, the structure of the spike protein of the Omicron variant consists of a higher percentage of alpha-helix structures, which are known for increasing conformational stability of proteins; therefore, we expect that its stability will also be increased (Slater, 2003; McKay et al., 2018; Poboinev et al., 2018). We predicted that T470-Q474 residues of the receptor-binding loop might undergo a coil to alpha-helix transition, which might facilitate the formation of strong binding with hACE2 and enhance transmissibility. However, there was no alteration in the disorder property of the spike protein in the Omicron variant than the reference, although the overall residual flexibility is reduced, which may affect its function. In contrast, flexibility was predicted to increase in the transmembrane domain of the S2 subunit of the spike protein, which might enhance viral fusion to the host cells, although conserved functional residues are predicted to present in less number (Römer et al., 2021). Contact map overlapping analysis supported the prediction that there are fewer conserved functional residues in the Omicron spike protein since we observed 90.5% common contacts with reference protein, suggesting differences among the functional residues. The receptor-binding domain of the proteins, however, showed 91.3% common contacts with an RMSD value of 1.07, and a TM-score of 0.96355, while the respective values for the whole protein were 0.2 and 0.99780. Based on these results, it is evident that the greater part of the changes took place in the RBD region, which can also be explained by the presence of half the mutations there. The functional RBD domain is essential for recognition of and binding to ACE2, so any mutation in the RBD domain could negatively influence its function as well as its affinity for binding to the ACE2 receptor (Scialo et al., 2020). The findings of our research suggest that three mutations in the RBD region could affect its functionality; fourteen mutations could increase the disease severity, while all mutations would decrease its stability.

Furthermore, a number of mutations in the Omicron variant have previously been reported in other VOCs, and all of these mutations correlated with immune evasion, increased transmissibility, and stronger binding to hACE2, so higher transmissibility and increased immune evasion are expected from the Omicron variant (Choi and Smith, 2021; Farinholt et al., 2021; Tatsi et al., 2021; Lindstrøm et al., 2022). The extent of viral transmissibility is largely determined by its ability to bind to its receptor, and we found that the Omicron variant binds to hACE2 more strongly than the reference strain. T478K and N501Y mutations have been reported to enhance the interaction with hACE2 in the Delta variant, which are also present in this variant. In RBD-hACE2 interactions, N501 together with Q493, Q498 residues are crucial because they form polar contacts with the ACE2 residues K31 and K353. The Q498R mutation caused a dramatic change in the protein-protein interactions network, as it formed several hydrogen bonds with V209, D216, E208 and Q210 residues, and ion interactions with E208 and D216 residues of hACE2. Moreover, the Omicron variant carries the D614G and P681H mutations, which have been previously described as critical mutations that enhance the transmission and infectivity of SARS-CoV-2 (Zhang et al., 2020; Lista et al., 2021; Zhou et al., 2021). Overall, we have observed, using the 6M0J PDB as reference structure that the number of interfacial contacts in the Omicron variant is significantly increased and that most are hydrogen bonds, which explains the increased binding affinity.

Lastly, the molecular dynamics simulations have learnt that residues Q474, G476, N477, K478, A484, F486, and Y489 at the interface enhanced the interactions by forming salt bridges and hydrogen bonds while maintaining the previous interactions. Additionally, on average, we have discovered 6.89±1.28 hydrogen bonds and -98.02 kcal/mol binding energy for the RBD of Omicron variant when bound to hACE2 while these values are 5.52±1.26 and -83.7 kcal/mol, respectively for the reference variant, which indicates better interaction of Omicron RBD with hACE2. A large number of mutations in the receptor-binding domain of spike protein increased the binding affinity with hACE2 and stabilized the RBD-hACE2 complex. For the Omicron hACE2-RBD complex, the RMSD and RMSF values were 2.05 Å and 1.7 Å, respectively, whereas, for the reference hACE2-RBD complex, they were 2.61 Å and 2.2 Å, which demonstrate fewer fluctuations among the atoms while bound with hACE2 than with reference spike protein. Therefore, molecular dynamics simulation data indicates the Omicron RBD-ACE2 complex is more stable with better binding energy than the reference variant.

## Conclusion

In light of the large number of mutations observed in the spike protein of the SARS-CoV-2, Omicron, a new variant of the virus poses serious concerns. Therefore, in the following work, we used *in silico* computational methods to study the dynamic changes in protein structure due to the mutations caused in spike protein. The results of our analysis revealed significant differences with respect to physicochemical, structural and functional changes, the enhanced binding affinity of RBD with hACE2, and a lower residual fluctuation, which might facilitate the greater transmission. The preliminary data about the rapid spread of this variant in different parts of the world is consistent with our observations. Consequently, we would like to point out that if the current trend of high viral transmissibility with the Omicron variant continues, it could become the predominant strain and cause yet another outbreak of COVID-19. Therefore, this variant should be subjected to more comprehensive research.

